# A Virus-Packageable CRISPR System Identifies Host Dependency Factors Across Multiple HIV-1 Strains

**DOI:** 10.1101/2022.11.03.515041

**Authors:** Vanessa R. Montoya, Trine M. Ready, Abby Felton, Sydney R. Fine, Molly OhAinle, Michael Emerman

**Author notes:** Corresponding author: Michael Emerman.

## Abstract

At each stage of the HIV life cycle, host cellular proteins are hijacked by the virus to establish and enhance infection. We adapted the virus packageable HIV-CRISPR screening technology at a genome-wide scale to comprehensively identify host factors that affect HIV replication in a human T cell line. Using a smaller, targeted HIV Dependency Factor (HIVDEP) sub-library, we then performed screens across multiple HIV strains representing different clades and with different biological properties to define which T cell host factors are strain-specific versus which ones are important across all HIV strains and different clades. Nearly 90% genes selected across multiple host pathways validated in subsequent assays as bona fide host dependency factors including numerous proteins not previously reported to play role in HIV biology such as UBE2M, MBNL1, FBXW7, PELP1, SLC39A7, and others. Our ranked list of screen hits across multiple viral strains form a resource of HIV dependency factors for future investigation of host proteins involved in HIV biology.

## Introduction

With a small genome of ∼9.2 kb that encodes 14 major proteins, HIV must hijack host cellular machinery to successfully establish infection. These host proteins, called dependency factors, are needed for entry, transit into the nucleus, uncoating, viral integration, as well as subsequent steps of transcription of viral RNA, RNA splicing and export, protein translation, and virion assembly and budding.

While many of these important host factors have been identified one by one, high throughput methods have also attempted to create comprehensive maps of host dependency and/or restriction factors. For example, four genome-wide RNA-interference screens (1-4) as well as a genome-wide CRISPR/Cas9 screen (5) sought to identify dependency factors for HIV infection. While these screens identified genes important for cellular attachment and entry, nuclear entry, integration, transcription, and nuclear export (1-5), there was very poor overlap across screens (6). Other approaches have sought to identify novel dependency factors through protein-protein interaction screens and identify hundreds human proteins that are physical interactors with the 18 HIV-1 proteins and polyproteins in Jurkat T-lymphocyte and 293T cells (7). A subset of these interacting proteins were functionally validated in primary CD4+ T cells as HIV dependency or restriction factors (8). In addition, interaction based and gene network approaches have been used to computationally predict dependency factor genes (9) and to develop a viral-host dependency epistasis map (Ve-MAP) (10). However, none of these approaches have used an assay with a functional readout to examine multiple viral strains across the entire HIV lifecycle in T cells.

We had previously developed a high throughput CRISPR screening method, called HIV-CRISPR, in which lentiviral genomes encoding sgRNAs are incorporated into budding virions serving as a readout for genes important for HIV infection (11-13). Although this work focused on finding restriction factors against HIV-1, we also demonstrated that this method could also identify dependency factors, although as the guide library used in those studies specifically targeted interferon-stimulated genes, the number of dependency factors identified was relatively limited. Here, we use a whole genome guide library to look more globally at HIV dependency factors in a T cell line. We used these data to inform the design of a smaller, custom HIV Dependency Factor CRISPR sublibrary (HIVDEP) to identify HIV dependency factors across multiple HIV strains. Our screens identified many previously reported HIV dependency factors across multiple parts of the viral life cycle, including the HIV-1 receptor CD4, co-receptor CXCR4, LEDGF/p75, NFkB, and many genes encoding components of the Mediator complex. Further, we identify genes not previously reported to play a role in HIV biology involved in transcriptional regulation, protein degradation, RNA regulation, as well as epigenetic factors affecting both early and late events of viral replication. Genes not previously identified as being important for HIV replication that were validated here include genes involved in Cullin-ring ligase mediated protein degradation, such as UBE2M and FBXW7, pre-mRNA alternative splicing regulator MBNL1, transcription factor PELP1, and zinc transporter/ tyrosine kinase activator SLC39A7. Our expanded catalog of ∼200 host dependency factors required at different stages of the life cycle has potential to inform therapeutic and cure design.

## Results

### A whole genome CRISPR screen for HIV dependency genes

The HIV-CRISPR screening method has previously identified restriction factors in THP-1 cells (a CD4+ monocytic leukemia cell line) that were involved in the interferon response against HIV using a focused CRISPR guide library of interferon-stimulated genes (ISGs) (11). In order to more comprehensively identify HIV dependency factors, we adapted this approach to a whole genome strategy utilizing the Toronto Knockout version 3 whole genome (TKOv3) library. This library targets 18,053 protein-coding genes with 4 guides per gene and also includes 142 non-targeting control guides (14).

Our screening strategy is outlined in Figure 1A. The HIV-CRISPR vector is a lentiviral vector that contains the sequences for a guide RNA (from an sgRNA library) targeting a single gene, the Cas9 enzyme, a viral packaging signal (psi), and a repaired long terminal repeat region which allows transcription of a genomic RNA (11). In addition to transcribing Cas9 and the encoded sgRNA, cells that are transduced with the HIV-CRISPR vector are also capable of making mRNA corresponding to full-length HIV-CRISPR genomes after infection with HIV-1. Thus, upon infection with an HIV strain of interest, newly-produced virions will also encapsidate the HIV-CRISPR genome *in trans*. Therefore, if a dependency factor has been knocked out, these cells will support less productive infection and fewer HIV-CRISPR genomes containing the sgRNA specific for the dependency factor will be packaged and released from infected cells. This relative depletion of sgRNAs in the viral supernatant, as compared to the genomic DNA (13) is the readout in which we infer knockout of a dependency factor. The advantage of this system is that CRISPR guides can be evaluated in bulk rather than arrayed in single wells, which makes a whole genome approach feasible.

**Figure 1.**
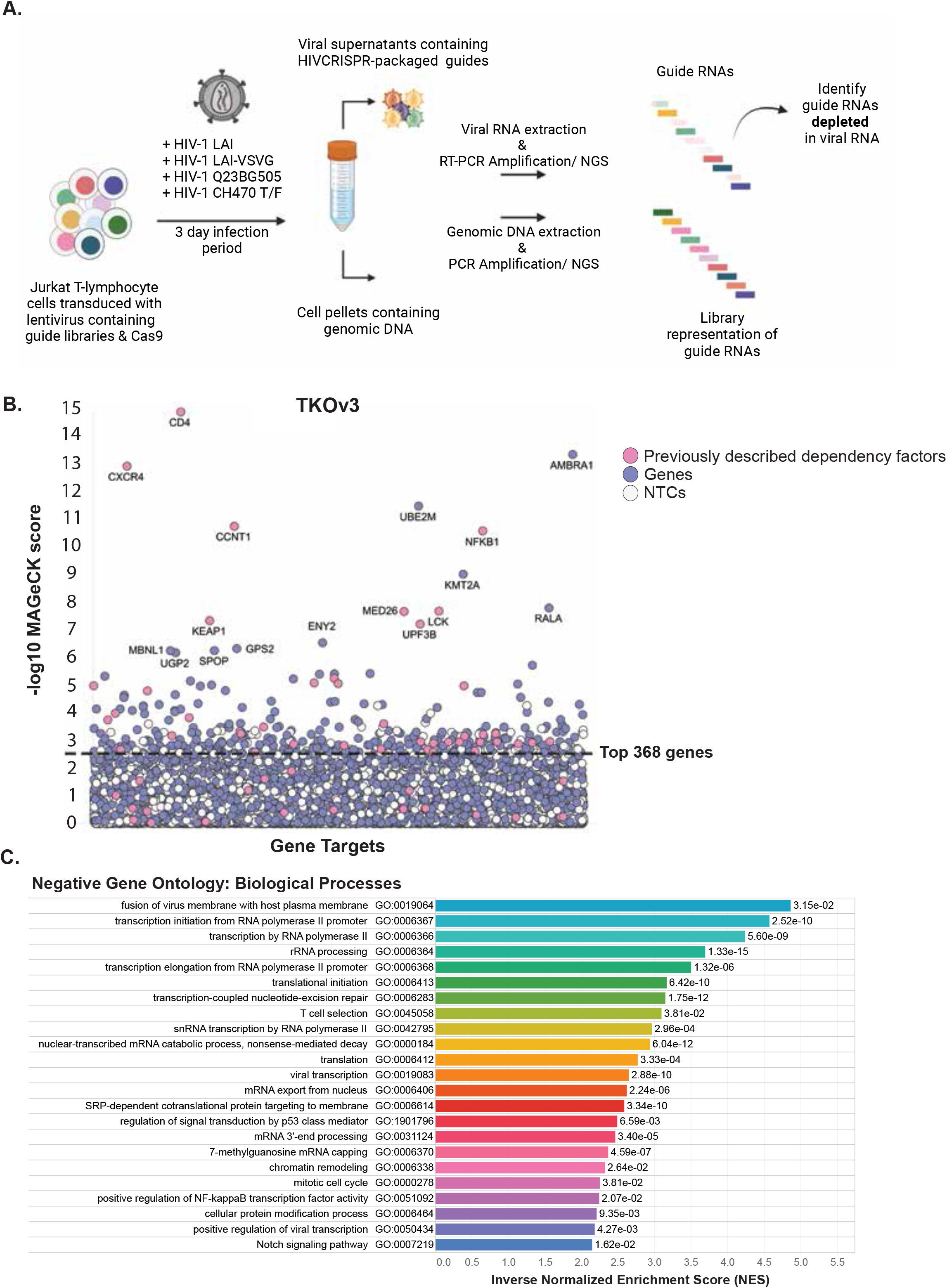
Genome-wide HIV-CRISPR screening to identify dependency factor candidates for HIV Dependency Factor guide library. **(A)** HIV-CRISPR screening process. Jurkat-CCR5 cells containing a CRISPR knockout library are infected with HIV-1 in duplicate infections. Strains used in this study are listed about the arrow. Viral RNA and genomic DNA were collected 3 days post infection and sequences corresponding to sgRNAs present in virions (vRNA) and genomic DNA (gDNA) were quantified through deep sequencing. **(B)** MAGeCK Gene analysis of the Genome-wide screen showing the most depleted (dependency) genes in Jurkat T cells infected with HIV-1_LAI_ in duplicate infections. <u>X-Axis</u>: randomly arrayed target genes. <u>Y-Axis</u>: -Log10 MAGeCK score for each gene. Host factors previously reported as dependency factors are shown in Pink. Other genes in the library are shown in Purple. “Synthetic” Non-targeting control (NTC) genes generated in silico by iterate random binning of the 142 NTC sgRNA sequences to generate a negative control set which are shown in White. Gene names are shown for all hits with a -Log10 MAGeCK score greater than 6. The top 368 most depleted candidate genes (excluding the synthetic NTC genes) are indicated by the dashed black line. **(C)** Gene Set Enrichment Analysis of the genome-wide screen. The top 20 most enriched Negative Gene Ontologies (most enriched pathways of the depleted/ dependency factor candidate genes) are shown here in ranked order by Inverse Normalized Enrichment Score (NES). Adjusted p values are displayed next to each gene ontology.

We transduced Jurkat-CCR5 cells, a T cell line susceptible to HIV infection and engineered to express the CCR5 co-receptor, with the HIV-CRISPR vector containing the TKOv3 whole genome library. The cells were then infected with HIV-1 at an MOI of 0.5, and genomic DNA and viral RNA was harvested from cell pellets and viral supernatants, respectively three days after infection. These samples were then deep sequenced to quantify enrichment or depletion of guide RNAs in the viral supernatants compared to guide representation in the cellular genomic DNA (13). Using the Model-based Analysis of Genome wide CRISPR/Cas9 Knockout (MAGeCK) screens algorithm and the log 2-fold change in gene depletion based on all four guide RNAs, statistical scores were assigned (15, 16) to delineate the most depleted genes, indicating factors important for infection (Figure 1B; Supplemental table 1).

**Table 1.** Viruses used in the HIVDEP screens reported in this study. LAI was used twice to assess reproducibility of the screen (referred to as “LAI redo” in this paper). Percent infection of each biological replicate of each HIVDEP screen (as measured through flow cytometry quantification of cytoplasmic gag 3 days post infection) is shown in the last column.

| HIV-1 strain | Clade | Co-receptor tropism | Traits | %p24 3dpi per biological replicate |
| --- | --- | --- | --- | --- |
| LAI | B | CXCR4 | Lab-adapted | LAI :<br>R1 = 10.57; R2 = 10.33 |
|  |  |  |  | LAI redo :<br>R1 = 7.05; R2 = 7.5 |
| LAIΔenv-VSV-G | B | n/a | Lab-adapted/ alternate entry mechanism | R1 = 23.93<br>R2 = 23.9 |
| Q23.BG505 | A | CCR5 | Primary isolate | R1 = 6.19<br>R2 = 6.32 |
| CH470 TF | B | CCR5 | Primary isolate/<br>Transmitted founder virus | R1 = 20.6<br>R2 = 22.9 |

As expected given their essential role in viral entry, the receptor CD4 and the co-receptor CXCR4 (required for the CXCR4-tropic strain HIV-1_LAI_ used in this screen), were among the most depleted genes, ranking as the top scoring dependency factors (Figure 1B). Additionally, many other previously reported dependency factors were identified in the 300 top scoring dependency factors (shown as pink dots) that include all different parts of the HIV lifecycle including entry (CD4, CXCR4, LCK), integration and uncoating (PSIP1, TNPO1), transcription (NFKB1, SP1, CyclinT1 (CCNT1)) Mediator complex genes (MED7, MED10, MED16, MED18, MED23, MED26), POLR2A), and budding (TSG101, CHMP4B). Importantly, the non-targeting controls randomly binned to synthetic “genes” cluster near the bottom.

As a global approach to identify the most negatively enriched biological processes of the genome wide screen (candidate dependency factor pathways), we used the Gene Set Enrichment Analysis as part of the MAGeCK-Flute pipeline (15). Of the top 20 negatively enriched Gene Ontology pathways (Figure 1C), three are classified as important for viral processes, including fusion of virus membrane with host plasma membrane (GO:0019064), viral transcription (GO:0019082), and positive regulation of viral transcription (GO:0050434). Notably, there were enriched pathways identified that may play roles for multiple stages of the viral life cycle, including entry/egress, transcription, mRNA capping and processing, translation, protein modification/trafficking, and T cell selection. Notwithstanding the caveat that the screen missed some expected dependency factors that did not cluster away from the NTCs (pink dots below the line in Figure 1B), this screening approach is capable of identifying dependency factors across the entire viral life-cycle and provides ample opportunity to investigate novel gene roles in the HIV life cycle.

### An HIV-dependency sub-library screened with multiple HIV-1 strains

Smaller, targeted libraries are more powerful as it is easier to maintain suitable coverage, more guides can be utilized per gene, and the smaller screens can be done in much higher throughput in more replicates. Therefore, before doing any validation of the hits in the whole genome screen, we used data from our whole genome to inform the construction of a targeted CRISPR guide library called the HIVDEP (HIV Dependency Factor) library. We initiated this library by including guides targeting genes in the top scoring 500 from the TKOv3 genome-wide screen. After subtracting non-targeting controls from the rankings, this amounted to 368 genes. We also added back the known dependency factors that fell below this threshold in the screen including CCR5 and PAPSS1 which are essential for CCR5-dependent strains. Because transcription and chromatin modification/ regulation were major categories of negatively enriched genes in the whole genome screen (Figure 1C), we also performed an additional targeted screen using a custom-designed Human Epigenome (HuEpi) library consisting of 838 human epigenome and epigenetic regulator genes (17) and included any gene with <10% false discovery rate (FDR) from the HuEpi screen that were not already included in the top TKOv3 gene list (Supplemental Figure S1A) as well as smaller number of negatively enriched genes in an ISG-related library (11) (Supplemental Figure S1B), resulting in an additional 131 genes. Finally, 11 genes previously called “non-essential” (18) that were neither depleted nor enriched in our genome-wide screen were included as negative control genes. In total, the HIVDEP library consists of 525 genes targeted by 8 guides and 210 NTC guides for a total of 4,401 guides (Supplemental Table 2).

To identify specific host factors that are either important for all HIV-1 strains, or ones that are strain-specific, we used the HIVDEP library screened against genetically distinct HIV-1 strains (Table 1). Specifically, these include HIV-1 LAI, which is in Clade B and uses co-receptor CXCR4, HIV-1 Q23.BG505 which is a clade A CCR5-tropic strain (19), and a transmitted/ founder virus from clade B that is R5-tropic, HIV-1 CH470 T/F (20). We also included an HIV-1 strain (LAI) that was deleted for its own envelope gene and pseudotyped with VSV-G to delineate entry-specific factors. Thus, we used HIV-1 strains that represent two clades (A and B), utilize different entry mechanisms, (-X4, -R5, or VSV-G), and are either lab-adapted vs. primary strains. Each screen was done in duplicate and the number of infected cells in each screen ranged from 6% to 23% (Table 1). In addition, we also performed screens with the HIV-1 LAI strain on two different occasions each with a separate transduction of the library followed by infection with different stocks of HIV-1 LAI (screens referred to as LAI and LAI redo in Figure 2).

**Figure 2.**
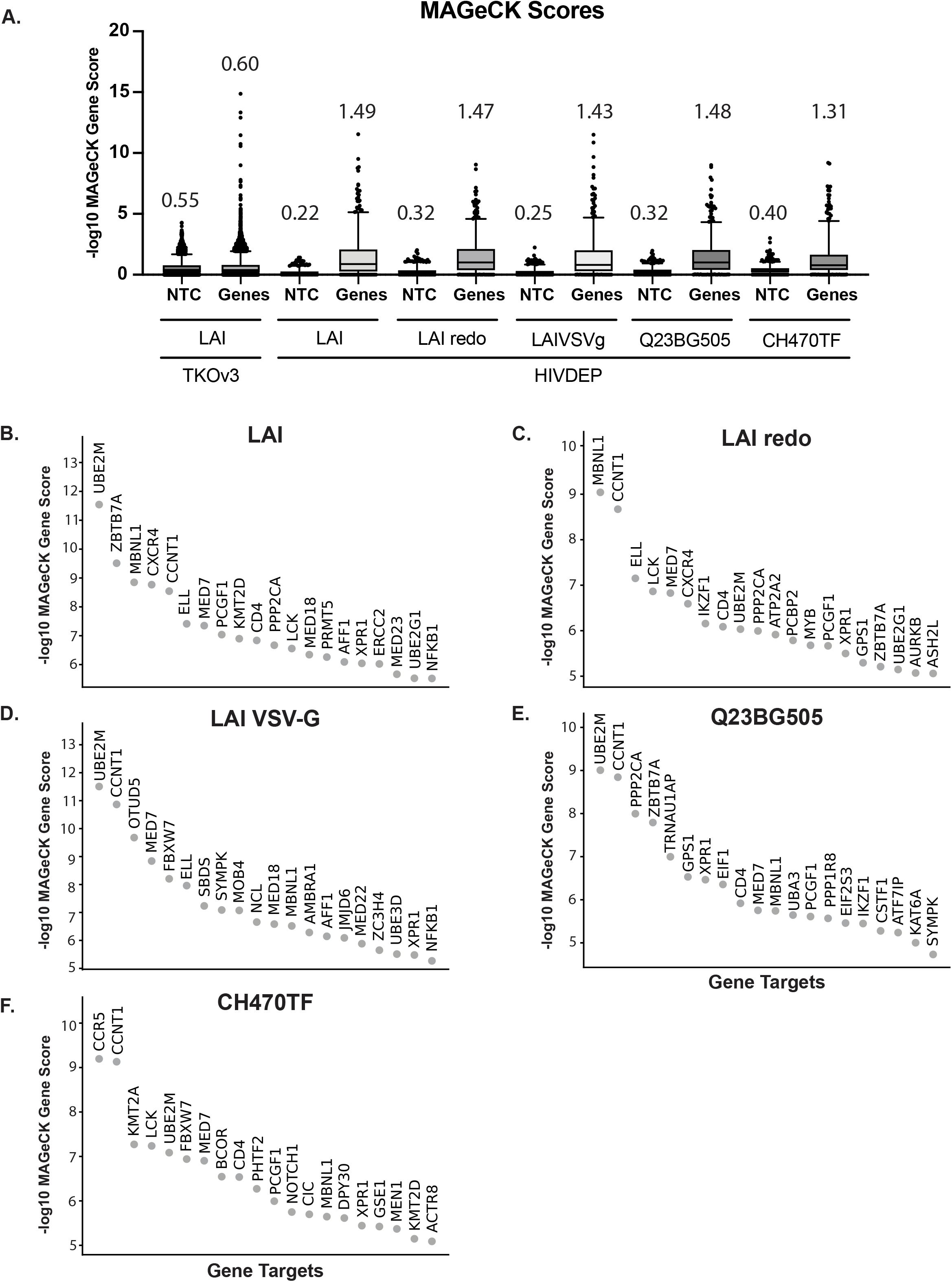
Iterative screening with HIVDEP sublibrary enriches for previously reported and candidate dependency factors. **(A)** MAGeCK score comparison of each CRISPR screen. Non-targeting control (NTC) sgRNA scores were randomly binned (four NTC guides per gene for TKOv3 or 8 NTC guides per gene for HIVDEP) to recapitulate the same number of genes in the respective libraries. <u>Y-Axis</u>: - Log10 MAGeCK scores. The mean MAGeCK scores for the synthetic NTC versus the genes are shown above each graph and are represented by the line within each box. The top 95^th^ percentile of NTC/genes is represented as the top horizontal line. For statistical analyses, the MAGeCK scores of the synthetic non-targeting controls (shown here as NTCs) were compared to the MAGeCK scores of the Genes in each screen. The Gene MAGeCK scores per each screen were also compared across screens. Each comparison resulted in significant p values =<0.0001 = ****; Welch’s t-test. Waterfall plots of the top 20 genes or all genes in each HIVDEP screen in descending order are shown for the following HIV-1 strains **(B)** LAI, **(C)** LAIredo, **(D)** LAI-VSV-G, **(E)** Q23BG505, and **(F)** CH470TF.

We compared the MAGeCK scores of genes versus the synthetic non-targeting controls (NTC) in the genome-wide screen versus each of the screens done with the dependency factor-focused HIVDEP library using box and whisker plots (Figure 2A). While the mean score of the synthetic non-targeting controls in each screen are similar, the mean of the MAGeCK scores for the genes in each HIVDEP screen was significantly higher than the mean of the genes in the genome-wide TKOv3 screen (Figure 2A; Welch’s t test). Moreover, the scores of the 95th percentile of genes (top horizontal line in each plot) were also increased in the HIVDEP library relative to the whole-genome screen for each of the HIV strains tested (Figure 2A). This indicates that the sub-library approach was successful in increasing the distance of noise to signal (NTCs to genes) relative to the whole genome screen. Moreover, waterfall plots of the top 20 hits from each screen (Figure 2B-2F) show that known dependency factors are identified such as the receptor CD4, CCNT1 (CycT1), NFKB1, and members of the Mediator complex. Moreover, several novel genes score in the top 20 for multiple strains, including UBE2M, MBNL1, FBXW7, PCGF1, and PPP2CA, implicating those gene products in processes important for viral infection across HIV-1 strains.

#### Validation of hits from the combined HIV dependency factor screens across different cellular processes

While hits across two screens done with the same strain are similar, the absolute MAGeCK scores between the two are not identical (LAI and LAIredo in Figure 2B and 2C), Therefore, to compare hits across screens, we normalized our data with a z score analysis as previously described (21). We thus created a ranked order list of both the average of all screen results by z score as well as the replicates from screens with each strain (Supplemental Table 3). We created pathway-focused heatmaps using the z scores for each replicate and the Gene Set Enrichment Analyses pathways established from the whole genome screen (Figure 3). As an initial control to determine if the screen could distinguish different strains, we looked at known entry factors that would be expected to be different between CXCR4-using, CCR5-using, and the VSV-G pseudotyped viruses in the Binding and Viral Entry Gene Set (Figure 3A). The HIV-1 LAI screens correctly identify CXCR4, but not CCR5 nor PAPSS1, while the opposite is true of the CCR5-tropic viruses, Q23BG505 and CH470TF. Each of the strains that uses wild-type HIV-1 envelope require CD4, but the VSV-G pseudotyped HIV-1 does not. LCK (lymphoid specific Src Kinase) which is important for T cell signaling downstream of CD4 engagement and reported to be important for viral core transit from the plasma membrane to the nucleus (22) tracks with CD4 in our screens (Figure 3A).

**Figure 3.**
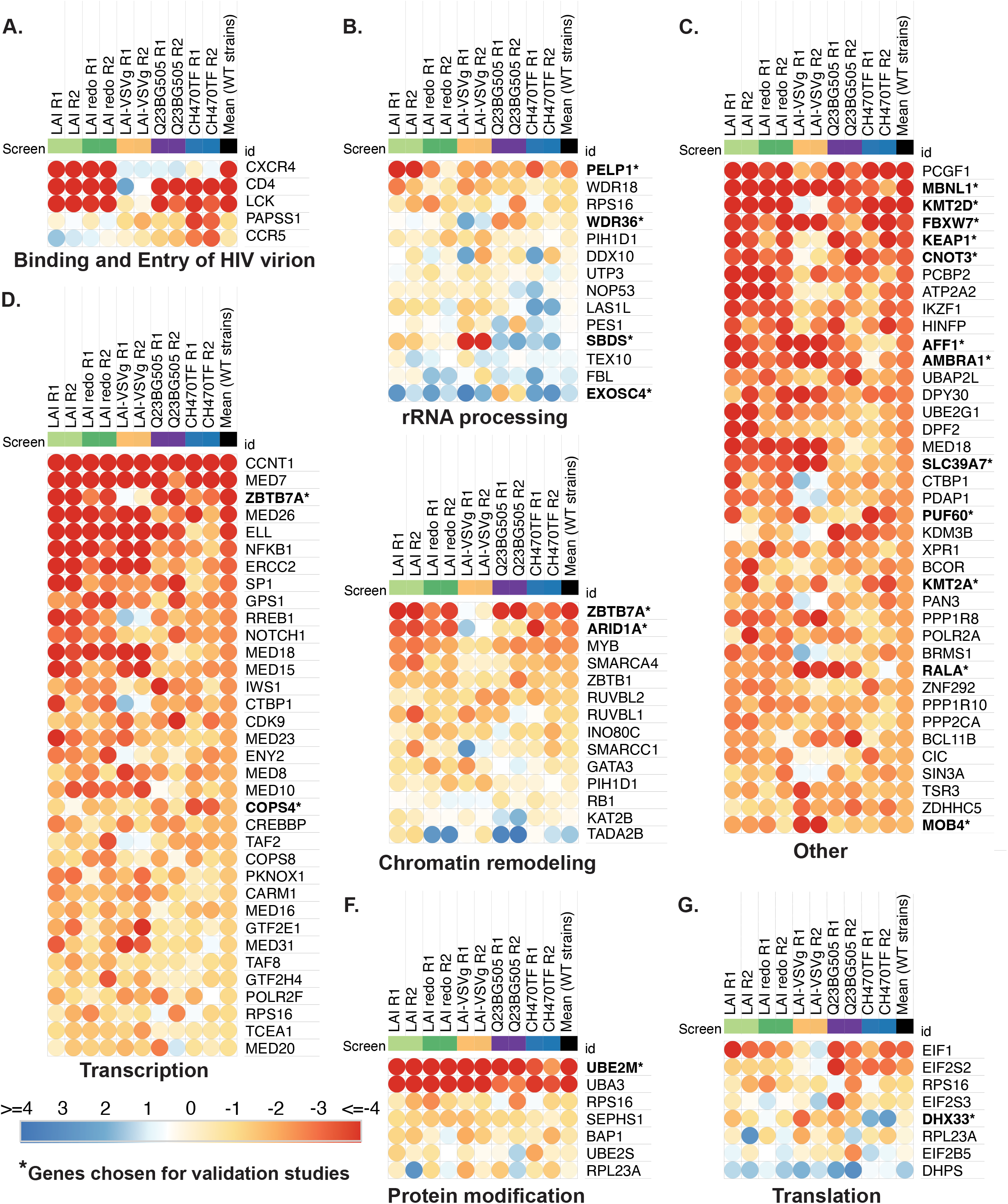
Common and differential use of host cellular pathways by HIV-1 strains. Comparative pathway-focused heatmaps showing enriched or depleted sgRNAs across each HIVDEP screen. The pathways shown are derived from the top 20 most enriched Negative Gene Ontologies of the genome-wide screen (Figure 1C). Any genes not included in the HIVDEP library were excluded. Z scores were calculated as described in (21). Z scores on each heatmap are colored from red (lowest/ most depleted genes, i.e. dependency factors) to blue (highest/ most enriched genes, i.e. negative or restriction factors). The median NTC z score was 0.5 and marks the inflection in the color scale. **(A-G)**. Each biological replicate is represented as a separate column showing the mean scores across wild type (non-VSV-G-pseudotyped) strains. The “Transcription” and “Other” heatmaps were truncated to the top 40 hits each heatmap. Gene names that are bolded with an asterisk indicate they were chosen for validation studies in Figures 4 and 5. “Other” are genes that were not assigned to any of the top Gene Ontology categories from Figure 1C.

The pathway-focused heatmaps also show enriched genes for each screen a large number for transcription as well as rRNA processing, protein modification, chromatin remodeling, and translation (Figure 3). We then chose a subset of genes across different host cellular processes (Figure 3) to functionally validate using infectious virus in multi-round spreading infections with both LAI and with Q23BG505. We also picked some genes that scored lower in the screens in order to determine a cut-off from which we would be more confident of remaining hits that were not directly validated, as well as some genes that scored more highly for one strain versus another (Figure 4A for the list ranked in descending order of average Z score as well as the Z score for each replicate of each gene). Our validation strategy used two guides per gene to knock out a candidate dependency factors in Jurkat-CCR5 cells. Positive controls included knocking out CD4, and negative controls included non-targeting control (NTC) guides, as well as guides targeting genes that are not involved in HIV or T cell biology, CD19 (B cell marker), and AAVS1 (the AAV integration site often used as a safe harbor locus (23-25)). The knockout cells were maintained as pools rather than clones to avoid clone-to-clone differences and were tested for gene editing as well as HIV infection within 2 weeks of generation.

**Figure 4.**
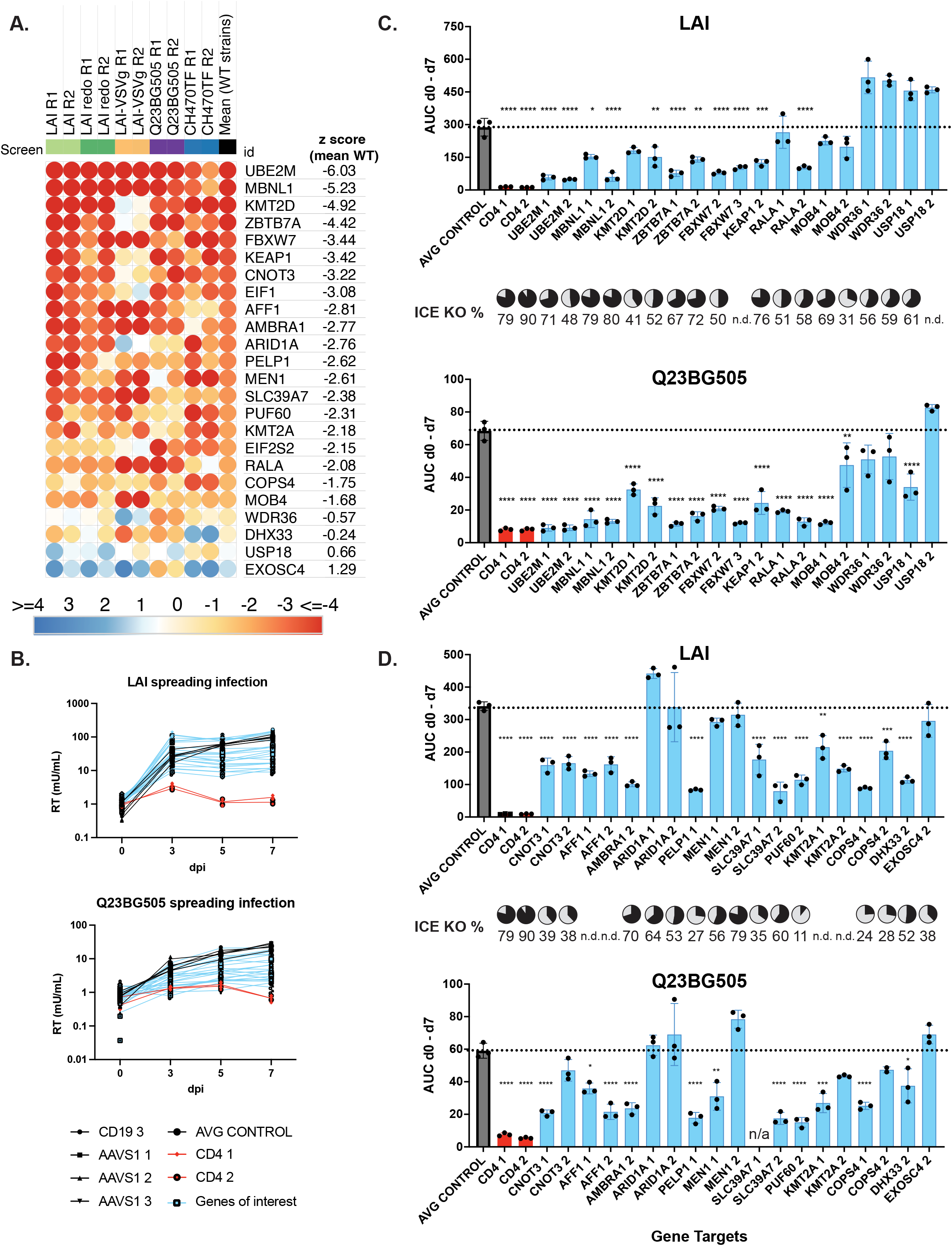
Validation of curated top hits list. (**A)** Heatmap of the candidate genes used for validation studies, ordered by the mean z score of WT strains **(B)** Pooled Knockout Jurkat-CCR5) generated by transducing with lentiviral vectors encoding sgRNAs including positive control gene CD4, negative controls CD19 or AAVS1, or candidate dependency factor genes. Two knockout lines per gene were generated using the highest scoring sgRNAs selected from across each HIVDEP screen. Viral supernatants were collected at days 0, 3, 5, and 7 to assess overall effect on replication kinetics via reverse transcriptase activity at each timepoint. The spreading infections were performed over two batches. <u>Y axis</u> = Reverse Transcriptase milliUnits/ mL. Batch 1 is shown in panel B and Batch 2 is shown in panel Supplemental figure S2 **(C and D)**. Area Under the Curve (AUC) was calculated for each cell line after 7 days of infection in Batch 1 (panel **C**) and Batch 2 (panel **D**) with either LAI or Q23BG505. Infection data of each guide is shown separately. For statistical analysis, all conditions are compared to the mean of the control cell lines (CD19 and AAVS1). One way Anova; Tukey’s multiple corrections test, p-value = ns > 0.05, <0.05 = *, =<0.05 = **, =<0.001 = ***, =<0.0001 = ****. For each knockout line, Synthego ICE analysis was performed and knockout scores are displayed as pie charts in line with the corresponding gene target. n.d. = could not be determined

Each knockout pool was infected with HIV-1 strains at an MOI of 0.15. All infections were done in triplicate and virus growth was measured by assaying reverse transcriptase activity released into the supernatant 0, 3, 5, and 7 days after infection. Replication curves for the majority of the genes of interest (blue) lines were in-between the CD4-KO lines and negative control lines (black), exhibiting inhibition of infection or slower growth kinetics for both LAI infection and Q23BG505 infections (Figure 4B and Supplemental Figure 2S). Infections were done in two batches with similar controls in both and grouped as such (Figure 4C is batch 1 and Figure 4D is batch 2). We used all of the data points in the spreading infection to calculate an area under the curve (AUC) for each infection (Figure 4C, 4D). We find that knockout of UBE2M which was one of the top hits in each of the screens (Figure 2B-2F) had the strongest phenotype in the spreading infections for both viruses tested (Figure 4C, 4D). In order of their effects, the strongest hits in this set of genes (aside from the positive control of CD4) were UBE2M, PELP1, MBNL1, ZBTB7A, and FBXW7. In addition, KMT2D, KMT2A, KEAP1, CNOT3, AFF1, AMBRA1, SLC39A7, PUF60, RALA, COPS4, and validated for at least for both viruses. MOB4 and WRD36 validated for Q23BG5050, but not for LAI. This was predicted for WRD36 since its z score was lower for Q23BG5050 (Figure 4A), although not expected for MOB4 which had similar z scores across the two viral strains in the HIVDEP screen (Figure 4A). We found similar results if we used the RT activity in the supernatant at day 5 or at day 7 as the primary readout instead (Supplemental Figure 3C, 3D), and find that at both time points, At the time of infection we also quantified the gene editing efficiency for each pool which were generally between 50% and 80% (pie charts in Figure 4). Therefore, as these are less than complete knockouts, the degree of decreased virus replication in the absence of the candidate genes is likely an underestimate. However, a lack of gene editing would not explain the failure of AIRID1A nor MEN1 to validate (Figure 4D). Nonetheless, these results show that about 90% of these selected screen hits validated as HIV dependency factors to some degree.

In order to test the hypothesis that the strength of effect of the knockouts on HIV infectivity is correlated to its z score in the screen, we graphed the z score against the respective amount of virus replication represented by the AUC (Figure 4C and D) in spreading infections. Indeed, there was a negative correlation between the strain specific z scores and the amount of virus replication produced in each knockout pool (Figure 5A); LAI (R^2^ = 0.39, p =0.001) and Q23BG505 (R^2^ = 0.41, p =0.001). We also find that this correlation holds if we use the mean z score of all wt strains tested in the screen ((Figure 5B); LAI (R^2^ =0.30, p = 0.007) and Q23BG505 (R^2^ = 0.46, p = 0.0004)) which argues that despite strain differences, the list of hits (Supplemental Table 3) can be used as a guide to identify HIV dependency factors. Thus, hits that scored higher in our screen are more likely to be authentic HIV dependency factors than those with lower z-scores.

**Figure 5.**
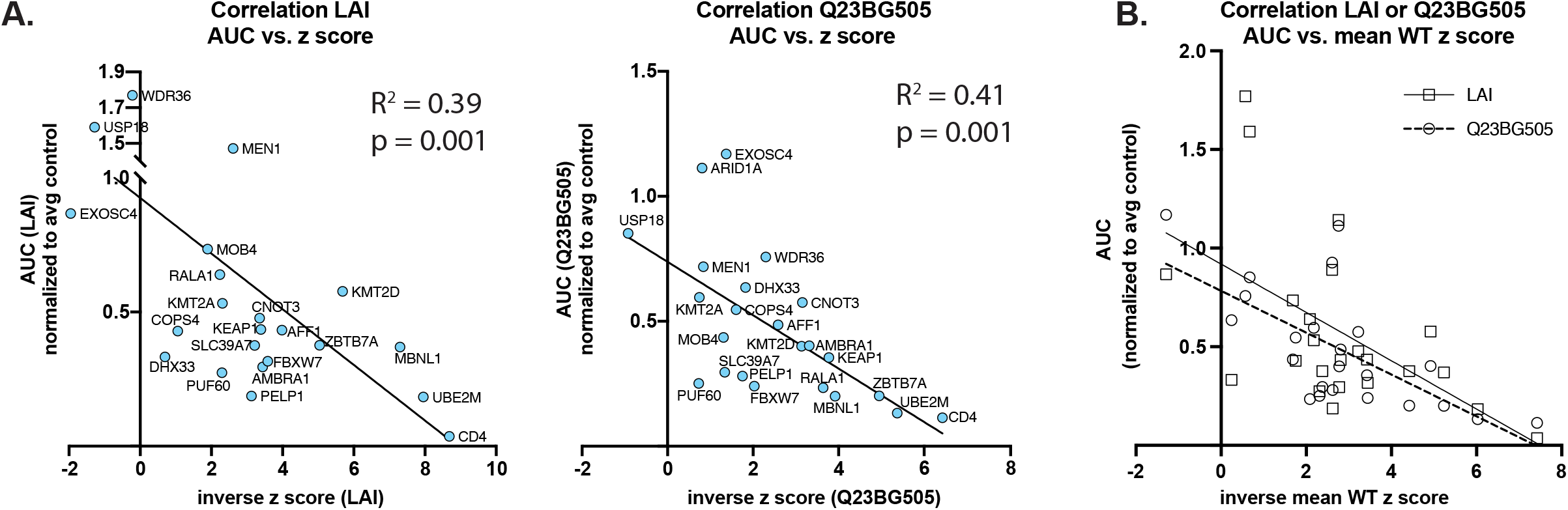
Correlation of gene z scores with AUC. **(A)** Area Under the Curve (AUC) was compared to the inverse z score of either LAI (left) or Q23BG505 (right). For statistical analysis, the mean biological replicate inverse z score per each gene for either LAI or Q23BG505, are compared to the mean AUC for infection of both guide knockouts per gene from Figure 4. Simple linear regression: LAI z score vs AUC (R^2^ = 0.39, p =0.001); Q23BG505 z score vs AUC (R^2^ = 0.41, p =0.001); **(B)** same as panel A, but the mean zscore for all wild-type strains (Supplemental Table 3) was used rather than strain-specific z score: Mean WT z score vs LAI AUC (R^2^ =0.30, p = 0.007); Mean WT inverse z score vs Q23BG505 AUC (R^2^ = 0.46, p = 0.0004). Squares and solid line represent LAI; circles and dashed line represent Q23BG505.

At the lower end of hits in our validation experiments, MOB4 (mean wt z-score -1.683) validated for Q23BG505, though not LAI while each gene with z-scores greater than MOB4 did not validate except for DHX33. WDR36, the next highest z-score gene we chose for validation, did validate for Q23BG505, but not for LAI, as predicted from the screen. We therefore set two tiers of z score cutoffs with different levels of stringency to best identify hits for future validation studies. The most conservative tier, Tier 1, was set to the MOB4 z score cutoff (Supplemental Table 3). The Tier 2 cutoff was set to the WDR36 z score. This is especially stringent as there are well studied and previously described dependency factors that score outside of Tier 1, but within Tier 2, including PSIP1/LEDGF (ranking #102 with mean wt z score of -1.37) which was also validated as a dependency factor in our system (Figure 6). This cutoff is likely still conservative as DHX33, which fell below Tier 2 (ranking #280), did validate for both viruses (Figure 4D), highlighting that there are likely other false negatives below this cutoff. We then conducted a literature review to identify how many of the 198 genes within Tiers 1 and 2 had already been identified in screens as HIV dependency factors as well as which ones have been experimentally validated (Supplemental Table 3). We find that 35 genes in this list had been previously identified in one or more previous screens (1-3, 5, 8). We find that 57 of the 198 genes in our Tier 1 and 2 groups had been validated as HIV dependency factors through functional studies. Therefore, we report here an estimated over 140 unstudied novel HIV dependency factors with more confidence in the hits with higher Z scores

**Figure 6.**
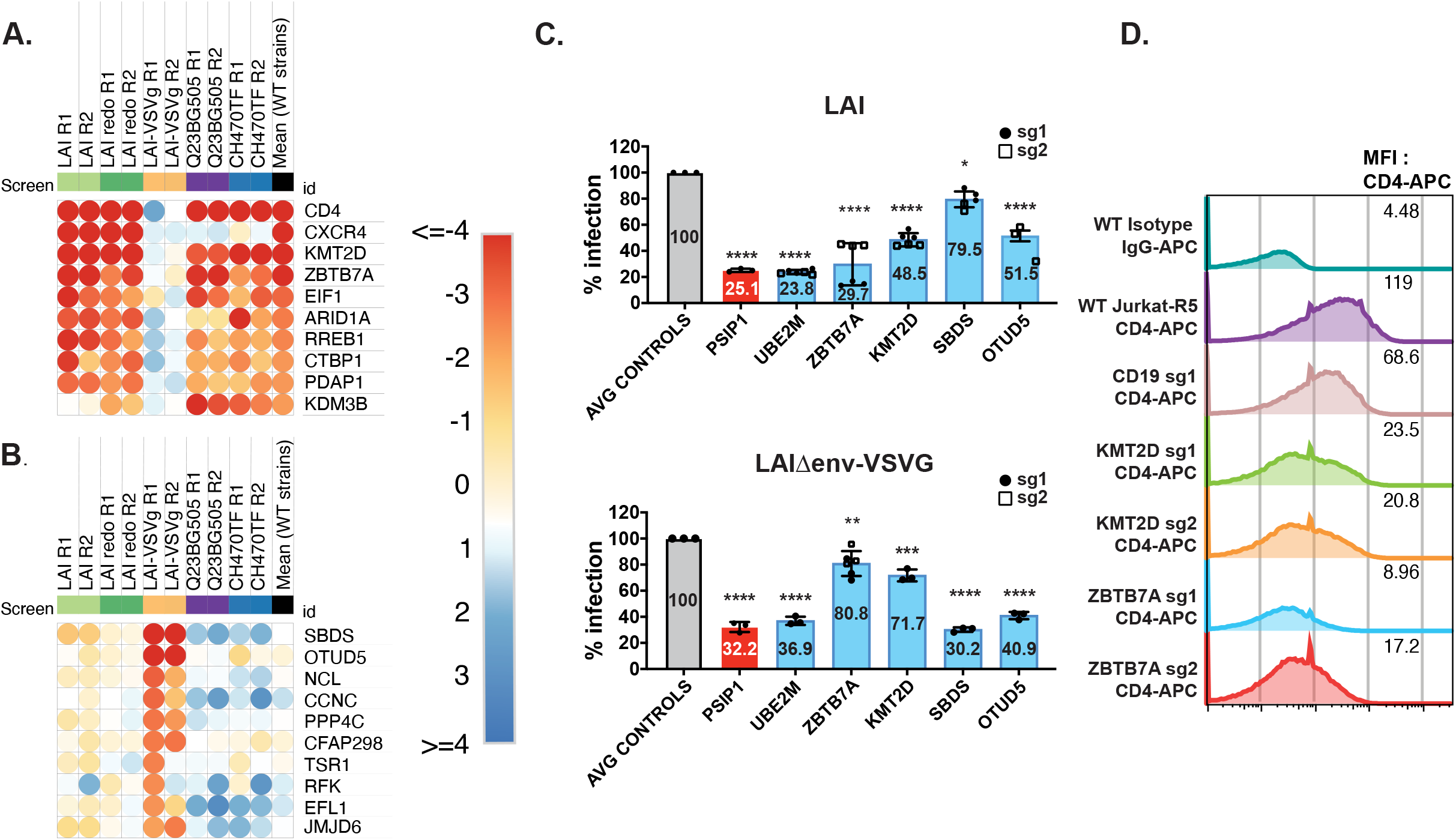
Entry-specific host factors. **(A)** Heatmap of the top 10 most depleted sgRNAs based on the mean of the wild-type strains, but not for VSV-G pseudotyped HIV-1. **(B)** Heatmap of the top 10 most depleted sgRNAs for VSV-G-pseudotyped HIV-1, but not for HIV-1 with wt HIV. Matrix for determination of panels A and B is in Supplemental Figure 3 with the heatmap z score values in Supplemental Table 3. **(C)** Jurkat -CCR5 cell pools edited for gene targets of interest were created by transducing wild-type Jurkat-CCR5 cells with lentiCRISPRv2 sgRNA constructs using sgRNAs for CD19 (B cell marker used as a negative control), PSIP1 encoding for p75/LEDGF (positive control), and genes of interest UBE2M, ZBTB7A, KMT2D, SBDS, and OTUD5), puromycin selected for at least 10 days. Knockout pools were infected with HIV 1-LAI or VSV-G-pseudotyped HIV-1 which both encode luciferase in place of the nef gene. Luciferase expression in infected wt or knockout cells was quantified 2 days post infection. All infections were done in triplicate using two pools of knockouts with one or two different sgRNA guides per gene. Data for the two different guides are shown as black circles (sg1) and open squares (sg2) for each pooled knockout cell line. The mean percent of luciferase activity of all replicates relative to the control cells is displayed on each bar. (**D)** Surface CD4 expression of WT or KO Jurkat-ZsGreen/CCR5 cells was quantified using flow cytometry for CD4-APC. MFI for each cell line is shown in each respective row. Two different knockout pools corresponding to two different guides per gene (with the exception of CD19) are shown.

### Combined HIV dependency screens highlight host entry factor preferences

As many experiments with HIV are done with VSV-G pseudotyped viruses to increase titers in single round infections, we were interested in further exploring hits with high differential z scores between the strains with wt HIV-1 envelope and VSV-G pseudotyped virus. Thus, we averaged the z scores for each gene across HIV-1 strains with wild-type envelopes and compared these scores with the z scores for the VSV-G pseudotyped virus (Supplemental Figure S3). Using this correlation matrix, we ranked the 10 most highly scoring genes scoring as dependency factors only for all HIV-1 strains with wt envelope but not LAI-VSV-G (Figure 6A) and separately ranked the 10 most highly scoring genes for VSV-G pseudotyped HIV but not for any of the wt HIV-1 stains (Figure 6B). We then functionally tested examples of each of these by knocking out the genes in Jurkat T cells and infecting them in a single round of infection with either an HIV-1 that either used VSV-G for entry or authentic HIV-1 envelope for entry. As positive controls, we used PSIP1 (also known as LEDGF), which should be needed for integration regardless of entry mechanism (26), and UBE2M which scored as one of the highest hits for all strains used in our screens regardless of entry mechanism or co-receptor (Figure 4). Negative controls included both Jurkat cells in which sgRNAs were used to target AAVS1, the safe harbor locus described above, as well as untransduced Jurkat cells.

We find that the decrease in infectivity of the cells knocked out for both PSIP1 and UBE2M was similar regardless of whether entry was through HIV-1 env or VSV-G (Figure 6C). Moreover, as predicted from the screen data, ZBTB7A and KMT2D were important for infection with HIV-1 envelope, but had a minimal effect on infection with the HIV-1 pseudotyped with VSV-G (Figure 6C). In contrast, knockout of SBDS affected the VSV-G pseudotyped virus, but not HIV-1 envelope-mediated entry while OTUD5 knockout had an effect on both viruses but decreased the VSV-G pseudotype to a greater extent. Examination of CD4 levels on the pools of knockout cells show that cells with ZBTB7A and KMT2D have reduced levels of cell surface CD4 relative to the control knockout cells (Fig. 6D). These results indicate some HIV dependency factors act indirectly on the virus by affecting receptor availability on the cell surface, and other factors that affect pseudotyped viruses may not be relevant for wt HIV. Thus, our screens with whole genome and more focused libraries identify both shared and strain-specific HIV dependency factors across a broad range of cellular processes.

## Discussion

We used HIV-CRISPR screening first at the genome-wide scale, and then at a smaller, targeted scale, to uncover novel genes that act as HIV-1 dependency factors. By using the HIV-CRISPR screening technique which assesses enrichment or depletion of guide-encoding genomes based on HIV-1 release from infected cells, we are able to establish a powerful functional assay to identify factors that promote HIV infection across the entire viral life cycle in a T lymphocyte cell line. Additionally, we were able to uncover dependency factors across multiple strains, from different clades and with different co-receptor tropisms. In addition to the genes above that were validated, we also conducted GSEA pathway analyses to investigate which host pathways to which HIV-1 is most dependent in these screens, with transcription-related pathways being the most represented.

### Transcription and chromatin remodeling factors as a major axis of HIV host dependency

Of the enriched pathways we identified, there were six transcription-related pathways (Figure 1C) which comprised a list of 58 genes that were included in the HIVDEP library as well an enriched pathway that includes chromatin remodeling genes. These are the largest pathway focused heatmaps, demonstrating an emphasis on transcription as a major focal point for host factors that promote HIV replication (Figure 3A). Some of the most enriched genes included previously reported dependency factors (positive controls) such as pTEFb components, CCNTI and CDK9, which are important for transcriptional elongation of HIV transcripts, (mean wt z scores: -7.97 and -2.18), transcription factors NFkB and SP1 (mean wt z scores: -3.24 and - 3.02), and several Mediator complex genes (MED7, MED26, MED18, MED15, MED23, MED8, MED10, MED16, MED20) (Supplemental Table 3). In addition to Tat-recruited pTEFb members, the Super Elongation Complex (SEC) has also been reported to be required for efficient Tat transactivation and elongation (27, 28). ELL is a member of the SEC and scored highly for all strains, except for CH470TF. Similarly, AFF1, which encodes for a scaffold protein of the SEC, also scored highly for each virus except CH470TF. AFF1 was also identified as a dependency factor in primary T cells (Hiatt et al. 2022). This is consistent with our data as we see a reduction of infection for both LAI and Q23BG505 in AFF1 KO Jurkat cells (Figure 4D). CREBBP, which encodes for CREB-binding protein (CBP), is recruited by Tat to the viral LTR (29), and acetylation of Tat by CBP/p300 has been shown to be important for transcriptional activation at the LTR (30, 31). Here, we have shown that CREBBP is important for four different viruses as it scored highly in our screens for LAI, LAI-VSV-G, Q23BG505, and CH470TF (mean zscore: - 1.767).

PELP1 was identified in an siRNA screen in HeLa-CD4 cells as a restriction factor (32). However, both our screens and validation experiments suggest that it is acts as a dependency factor in Jurkat cells (mean replicate wt z score: -2.617) (Figures 3B, 4D). PELP1 has been shown to interact with SETDB1, a methyltransferase, oncogene, and restriction factor that effectively inhibits Tat activity by methylation (33). It is possible that PELP1 promotion of SETDB1-Akt activation (34) may draw SETDB1 away from Tat, thus allowing transactivation. Therefore, PELP1 co-regulation of HIV-1 transcription factors and SETDB1 activity could explain the dramatic reduction we see upon gene knockout in Jurkat T cells.

ZBTB7A (also known as FBI-1 and LRF) is a zinc-finger protein involved in a diverse array of activities involving transcriptional co-repressors (35). It has also previously been identified as a protein that binds the HIV-1 LTR and associates with HIV-1 Tat (36, 37). Notably, ZBTB7A was recently identified to affect human coronavirus 229E through modulation of oxidative stress (38). However, we find that ZBTB7A was a hit for each virus except for the VSV-G pseudotyped virus (mean replicate z score of 0.2) and resulted in lower cell surface CD4 levels in the knockout pools implying ZBTB7A transcriptionally regulates pathways important for CD4 levels rather than HIV transcription itself. Similarly, KMT2D, also known as MLL2, is a lysine methyltransferase, but known for post-translational modification of histones reviewed in (39) appears to act indirectly on HIV through effects on CD4 levels.

### Cellular Protein Modification as HIV dependency factors

Our screens identified multiple genes involving cullin-mediated ubiquitin ligase complexes. Cullin ring ligases are multi-subunit E3 ubiquitin ligase complexes responsible for ubiquitylating ∼20% of cellular proteins targeted for degradation (40). Cullin ring ligases (CRLs) must undergo neddylation to be activated. Neddylation involves the transfer of ubiquitin-like molecule NEDD8 to a lysine residue of a substrate, subsequently affecting activity, conformation, and/or subcellular localization. Notably, UBE2M, which neddylates CRL-1, CRL-2, CRL-3, and CRL-4 via the substrate specific E3 ubiquitin ligase Rbx1 (mean wt zscore: -0.54), was one of the top scoring hits across all screens, including the genome-wide LAI screen and the HIVDEP LAI VSV-G screen, following essential dependency factors CD4 and CyclinT1 (mean wt zscore CCNT1: -7.97; mean wt zscore CD4: -7.42; mean wt zscore UBE2M: -6.03).

Knockout of UBE2M in Jurkat cells exhibited the most potent inhibition of infection of all the validated genes except for the positive control, CD4 (Figure 4). Neddylation and specifically UBE2M has been shown previously to be important for activation of CRL-4 and CRL-5 for efficient degradation of restriction factors SAMHD1 by HIV-2 Vpx and APOBEC3G by Vif, respectively (41). However, as neither of these factors are important in Jurkat T cells infected with wild type HIV, we hypothesize we have identified other effects of UBE2M neddylation for HIV-1 replication. UBA3 is one of two components of the NEDD8 E1 Activating Enzyme (NAE) heterodimer, just upstream of UBE2M in the neddylation process (42). UBA3 scored extremely well across all screens, including the LAI-VSV-G pseudotyped virus (Figure 3F, mean wt zscore: -4.258; mean zscore of all: -4.61). Notably, inhibitors targeting the NAE complex have been effective in inhibiting HIV-1 infection (41), further validating these degradation pathways as important for the virus to replicate.

FBXW7, a substrate receptor for degradation of proteins dependent on phosphorylation status has a large array of transcriptional effects in cells (43) and was also validated in our screens (Figure 3C; Figure 4) as an HIV dependency factor. FBXW7 has many substrates through which there are downstream effects. Interestingly, FBXW7 substrate CCNE2 (encodes for protein Cyclin E1), and Cdk2 and Cyclin E have previously been shown to induce HIV Tat activity by phosphorylation of the Ser16 residue, thus acting as a dependency factor (44-47). It is possible that HIV-1 hijacks FBXW7 to sequester it, thus preventing or co-opting degradation of CCNE2 or one of its other substrates that have no previous implication for HIV-1 infection.

### Additional HIV Dependency Factors

Of the genes chosen for validation studies, we validated 88% of these as dependency factors for at least one virus (Figure 4). Thus, as there are 72 genes in Tier 1 and 198 genes in Tier 2, if we use this percentage of validation rate as a rough estimate across the genes in Tier 1 and Tier 2, we estimate our CRISPR screen has, at minimum, identified at least more 156 HIV dependency factors in addition to the 18 validated here (Supplemental Table 3). However, this is likely a minimum number as there are known HIV dependency factors that are below #198 (Supplemental Table 3) on the list. For example, known dependency factors for budding TSG101 (ranking at #299 with mean wt z score = -0.12) and CHMP4B (ranking at #147 with mean wt z score = -0.91) did not meet this cutoff. Thus, the absence of a gene in our top list is not evidence of its lack of effect, but as we find a correlation between Z score and the magnitude of effect in spreading infections (Figure 6), in general, the higher the Z score, the more likely lack of that gene product will affect HIV replication. There are certain caveats to our screen results. First, the screen is done in Jurkat T cells, so that host genes that are more important in primary cells could be missed in our system.

There was only a small overlap between our HIVDEP guide library and the library of interactors that were validated in (8), however, we did identify some of the same genes (Supplemental Table 3). HIVDEP was also designed based on a genome-wide screen using LAI, a CXCR4-tropic strain and therefore some genes only important for another strain could also have been missed. Despite these limitations, we have identified and validated many new dependency factors for HIV-1, including factors that are important across clades, as well as many that are strain-specific. This list of host dependency factors can be used to further explore which pathways HIV-1 must hijack and has substantial implications for anti-HIV-1 therapeutic design and HIV cure strategies.

## Acknowledgements

We thank Nicholas Chesarino for critical feedback of this manuscript, Ferdinand Roesch for developing initial protocols for a whole genome screen with HIV-1, and all members of the Emerman lab for helpful suggestions and technical assistance. We thank Julie Overbaugh for sharing the pQ23.BG505 plasmid, John T. Poirier for advice and sharing code for z score analysis of CRISPR screen data, Andrew McOlash for executing the z score analysis code on our CRISPR datasets and data visualization advice, Matt Fitzgibbon, Qing Zhang, and Pritha Chanana at the FHCC for bioinformatics support, and Michael Zager for pipeline and data visualization advice and support. This work was supported by a Cellular and Molecular Biology Training Grant (T32 GM007270) awarded to V.M., NIH/NIAID P50AI150476 (principal investigator [PI], Nevan Krogan; subaward to M.E.), and NIH/NIAID R01 AI030927 to M.E. and NIH/NIAID R01 AI147877 to M.O. This research was supported by the Genomics, Bioinformatics, Data Visualization, and Flow Cytometry Shared Resources, RRID:SCR_022606, of the Fred Hutch/University of Washington Cancer Consortium (P30 CA015704).

## Materials and Methods

### Cell Culture

The Jurkat (ATCC) cell line was cultured in RPMI-1640 medium (Thermo Fisher Scientific) supplemented with 10% Fetal Bovine Serum (FBS), Penicillin-Streptomycin (Pen/Strep), and 10 mM HEPES. 293T and TZM-bl cells (ATCC) were cultured in DMEM (Thermo Fisher Scientific) with 10% FBS and Pen/Strep. Mycoplasma testing was done prior to screens and validation studies by the Fred Hutch Specimen Processing/Research Cell Bank Shared Resource and was not detected in any cell lines used in these experiments.

### Plasmids

The HIV-CRISPR vector was previously described (11). HIV-CRISPR constructs targeting genes of interest were cloned by annealing complementary oligos with overhangs that allow directional cloning into HIV-CRISPR using the BsmBI restriction sites (Supplemental Table 2). lentiCRISPRv2 plasmid was obtained from Feng Zhang via Addgene #52961. pMD2.G and psPAX2 were obtained from Didier Trono via Addgene #12259/12260. sgRNAs designed to target genes of interest were cloned into lentiCRISPRv2 using methods previously published (11). The wild type (HIV-1_LAI_) and *env*-deleted (HIV-1_LAI_VSV-G) HIV-1_LAI_ proviruses were previously described (48, 49). The Clade A HIV-1_Q23.BG505_ molecular clone was previously described (19, 50). CH470TF was provided by Beatrice Hahn (51). The lentiviral pHIV-dTomato (Addgene plasmid; 21374; RRID:Addgene_21374) and pHIV-Zsgreen (Addgene plasmid; 18121; RRID:Addgene_18121) expression vectors were deposited by Bryan Welm. The pHIV-zsGreen/CCR5 and pHIV-dTomato/CCR5 constructs were created by cloning the human CCR5 CDS into pHIV-zsGreen and pHIV-dTomato backbones using the BamHI and NotI sites. The Toronto human knockout pooled library (TKOv3) was obtained from Jason Moffat via Addgene #90294 (14).

### HIV Dependency Factor (HIVDEP) CRISPR/Cas9 sgRNA Library Construction

The HIVDEP sgRNA library is composed of 525 genes (4,191 sgRNAs). The top scoring 368 genes (as determined by -log10 MAGeCK score) of the genome-wide screen were included in the HIVDEP library. Additional unique top scoring genes with <10%FDR from HuEpi and PIKA screens were added to the HIVDEP library that contained an additional 131 genes. NR5A1, NHLRC4, SFTPA2, ZNF768, MYL10, GIMAP5, SPG21, CHSY3, ZNF25, REG1A, and ATXN3 were manually selected as non-essential genes (18) that were neither enriched nor depleted in the genome-wide screen. Six new sgRNAs were designed using two algorithms, GUIDES (52) and CHOPCHOP (53). We also manually included other known dependency factors, including CCR5 and PAPSS1. Two hundred and twelve Non-targeting controls were designed using GUIDES and also included. The HIVDEP sgRNA library was synthesized (Twist Biosciences) and cloned into HIV-CRISPR. Oligo pools were amplified using Herculase II Fusion DNA Polymerase (Agilent; 600677) combined with 1 ng of pooled oligo template, primers ArrayF and ArrayR (ArrayF primer: TAACTTGAAAGTATTTCGATTTCTTGGCTTTATATATCTTGTGGAAAGGACGAAACACCG and ArrayR primer: ACTTTTTCAAGTTGATAACGGACTAGCCTTATTTTAACTTGCTATTTC TAGCTCTAAAAC), an annealing temperature of 59°C, an extension time of 20 seconds, and 25 cycles. Following PCR amplification, a 140 bp amplicon was gel-purified and cloned into BsmBI (NEB; R0580) digested HIVCRISPR using Gibson Assembly (NEB; E2611S). Each Gibson reaction was carried out at 50°C for 60 min. Drop dialysis was performed on each Gibson reaction according to the manufacturer’s protocol using a Type-VS Millipore membrane (VSWP 02500). 5 μl of the reaction was used to transform 25 μl of Endura electrocompetent cells (Lucigen; 60242-2) according to the manufacturer’s protocol using a Gene Pulser (BioRad). To ensure adequate representation, sufficient parallel transformations were performed and plated onto carbenicillin containing LB agarose 245 mm x 245 mm plates (Thermo Fisher) at 492-times the total number of oligos of each library pool. After overnight growth at 37°C, colonies were scraped off, pelleted, and used for plasmid DNA preps using the Endotoxin-Free Nucleobond Plasmid Midiprep kit (Takara Bio; 740422.10). The HIVDEP library was sequenced and contains all 4,191 sgRNAs included in the synthesis (GEO Dataset, submission in progress).

### Virus and Lentivirus Production

293 T cells (ATCC; CRL-3216) were plated at 1.5×10^5^ cells/mL in 6 well plates one day prior to transfection. 3 ul of TransIT-LT1 reagent (Mirus Bio LLC) transfection agent was used per ug of DNA. For lentiviral preps, 293Ts were transfected with 667 ng lentiviral plasmid, 500 ng psPAX2 and 333 ng MD2G. For HIV-1 production, 293Ts were transfected with 1 ug/well proviral DNA. One day post-transfection media was replaced. Two- or three- days post-transfection lentiviral supernatants of the same type were combined and filtered through a 0.2 μm filter (Thermo Scientific; 720-1320). For lentiviral preps used for the creation of CRISPR/Cas9 knockout lines, supernatants were harvested and clarified from three wells per each lentiviral prep and were concentrated in microcentrifuge tubes for 1 hr at 4°C at 16,100 x g. The volume was reduced to equal 4x concentration before vortexing vigorously and resuspending at 4°C for 2 days. Each tube was combined per lentiviral stock before using for transduction. For HIVDEP library preps, supernatants from 40 × 6 well plates were pooled and concentrated by ultracentrifugation as described in (13). Concentrated lentivirus was used immediately or aliquots were made and stored at -80°C. To increase infectivity of the CH470 TF stock, we concentrated ∼75 mL of virus to ∼3 mL (25x concentration) using an Amicon Ultra-15 Centrifugal Filter Unit (Millipore Sigma; UFC905008). All viral and lentiviral infections and transductions were done in the presence of 20 μg/mL DEAE-Dextran (Sigma; D9885).

### HIV-CRISPR screening

Prior to the HIV-CRISPR screening or generation of knockout cell lines for validation studies, we performed CRISPR/Cas9-mediated knockout of Zinc Antiviral Protein ZC3HAV1 (ZAP) to increase efficiency of the HIV-CRISPR vector (11). We used gene Knockout v2 kit (GKOv2) for ZAP, including the following sgRNA sequences: GTGGTGTTGGAGACCGG, CCTGGAGCAGCGCGTCC, and TGAAGCAGCACACCTCC (Synthego, Redwood City, CA) complexed with 1 μL of 20 μM Cas9-NLS (UC Berkeley Macro Lab) and single cell sorted into a 96-well U-bottom plate to make clonal knockouts. Clonal KO lines were identified and selected using the ICE editing analysis software (Synthego). An individual clone with biallelic knockouts were used to create both ZsGreen/CCR5 and dTomato/CCR5 stably expressing lines by transduction with pHIV-ZsGreen/CCR5 or pHIV-dTomato/CCR5 lentiviruses followed by cell cytometry sorting for expression of CCR5 via ZsGreen or dTomato fluorescence using the BD FACSCANTO II or Sony MA900 (Fred Hutch Flow Cytometry Core) and analyzed with FlowJo software. These Jurkat cells were transduced with HIV-CRISPR vectors containing the TKOv3, HuEpi, PIKA, or HIVDEP library at an MOI of <1 and selected for puromycin resistance at 0.4 ug/mL. Ten days post puromycin selection, cells were infected with HIV-1 strains and levels of infection were measured by intracellular p24gag staining (Table 1) to obtain at least an MOI of 0.1. To maintain ∼500X coverage of sgRNAs for the HIV-1 Q23BG505 screen which was more difficult to achieve an MOI of 0.1, we increased the number of cells infected proportionally to the lower MOI (Table 1). Genomic DNA and viral RNA was harvested and amplified 3 days post infection, sequenced, and analyzed using the MAGeCK-FLUTE algorithm (15).

### Statistical analysis of HIV-CRISPR screen data

Library pools were demultiplexed, reads were assigned of libraries to assign sequences to each sample (allowing no mismatches), trimmed, and aligned to the TKOv3 or HIVDEP sgRNA libraries using Bowtie (54). An artificial or “Synthetic” NTC sgRNA set the same size of TKOv3 or HIVDEP was created by iteratively binning NTC sgRNA sequences (4 NTC sgRNAs/ gene or 8 NTC sgRNAs/gene for TKOv3 or HIVDEP, respectively). Relative enrichment of sgRNAs and genes were analyzed using the MAGeCK-Flute statistical package (15). Enriched gene ontologies of screen data were determined by Gene Set Enrichment Analyses (55) as a part of the MAGeCK-Flute package. For each biological HIVDEP screen replicate, z scores were calculated for each gene for comparison across screens performed with different viruses at different points in time. These were calculated as is described in (21). These z scores and enriched pathway data of the TKOv3 screen were used to generate pathway-focused heatmaps across each HIVDEP screen using the Morpheus (https://software.broadinstitute.org/morpheus).

### Dependency factor validation through spreading infections and luciferase quantification

Twenty-four candidate genes were knocked out in Jurkat-CCR5 cells using the two most efficient guides per gene from the sgRNA library (as calculated by log_2_ fold change sgRNA enrichment). We also generated non-targeting control (NTC), CD19-, AAVS1-, and CD4-CRISPR/Cas9 knockout Jurkat-CCR5 cells as negative and positive controls. Knockout cell pools were created via transduction with lentiCRISPRv2 containing gene targeting constructs. Less than 24 hours post transduction, media was replaced with RPMI containing 0.4 ug/mL Puromycin. Transduced cell pools were selected with puromycin for 10 days prior to HIV infection. gDNA was harvested for editing analysis after 10-12 days under selection. Knock out cells were maintained as pools rather than individual clones to remove artifacts of clone to clone heterogeneity in infection. We measured infection of the knockout pools by both luciferase luminescence at two days post infection and by measuring virus release three days after infection by measuring reverse transcriptase activity in viral supernatants as described (56). A stock of HIV-1 LAI was titered, aliquoted at −80°C and used as the standard curve in all assays.

### Genomic Editing Analysis

Knockout cells were harvested and genomic DNA was extracted using the QIAamp DNA Blood Mini Kit (Qiagen; 51185). Sites of editing were amplified using primers specific to each targeted locus (Supplemental Table 3). Primers were designed to amplify a 500 or 1000 base pair amplicon of the targeted locus using either Q5 High-Fidelity DNA Polymerase (NEB; M0491S) or Platinum Taq DNA Polymerase High Fidelity (ThermoFisher Scientific; 11304011). PCR amplicons were sequenced (Fred Hutch Shared Resources Genomics Core – sanger sequencing) and analyzed by ICE (Synthego) to determine the percent of alleles edited at each locus in the cell population (57).

### Flow Cytometry/p24gag Analysis

Intracellular p24gag staining was conducted on cells 3 days post infection to determine viral titer of each stock before using these for CRISPR screens or infection experiments with LAI, LAI-VSV-G, Q23BG505, and CH470TF. Cells were harvested and fixed in 4% paraformaldehyde for 10 min and diluted to 1% in DPBS. Cells were permeabilized in 0.5% Triton-X for 10 min and stained with 1:300 KC57-FITC (Beckman Coulter 6604665; RRID: AB_1575987) or 1:300 KC57-RD1 (numbers). Cells were analyzed on a Celesta or Fortessa Flow Cytometer (Fred Hutch Flow Cytometry Core). To assess infectivity of viral stocks used for the spreading infections, cells were fixed in BD Fix/Perm for 20 minutes, washed, and permeabilized in in BD Perm/Wash Buffer. The cells were stained with 1:300 KC57-FITC (Beckman Coulter 6604665; RRID: AB_1575987) or 1:300 KC57-RD1 (Beckman Coulter 6604667 in BD Perm/Wash Buffer for 30 minutes, washed, and then resuspended in PBS. Cells were read on a Celesta 2 or 3 (Fred Hutch Flow Cytometry Core) and analyzed in FlowJo. For CD4 cell surface marker staining, cells were washed twice in PBS, stained in PBS/1% BSA, incubated at 4°C for 1 hr, washed twice in PBS, and resuspended in PBS. The cells were stained with 1:50 CD4-APC (BD Biosciences 555349; AB_398593) and analyzed on Celesta 2 flow cytometer (Fred Hutch Flow Cytometry Core).

