## Supplemental Figures for "A Virus-Packageable CRISPR System Identifies Host Dependency Factors Across Multiple HIV-1 Strains"

#### Supplemental Figure 1

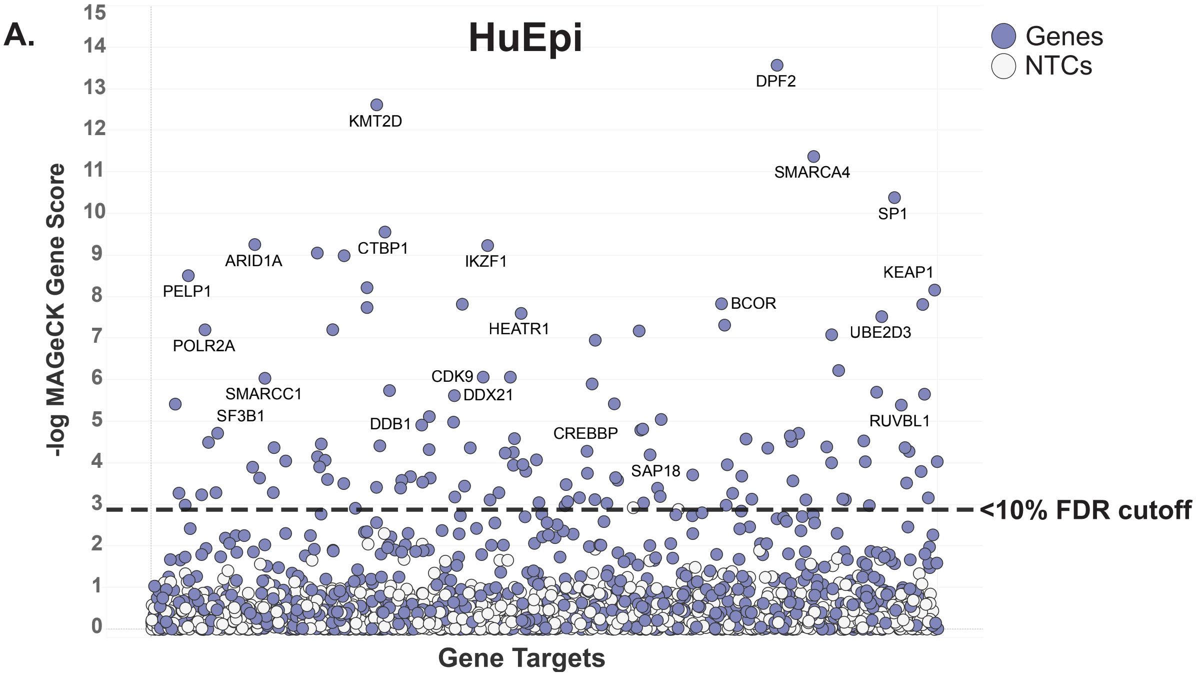


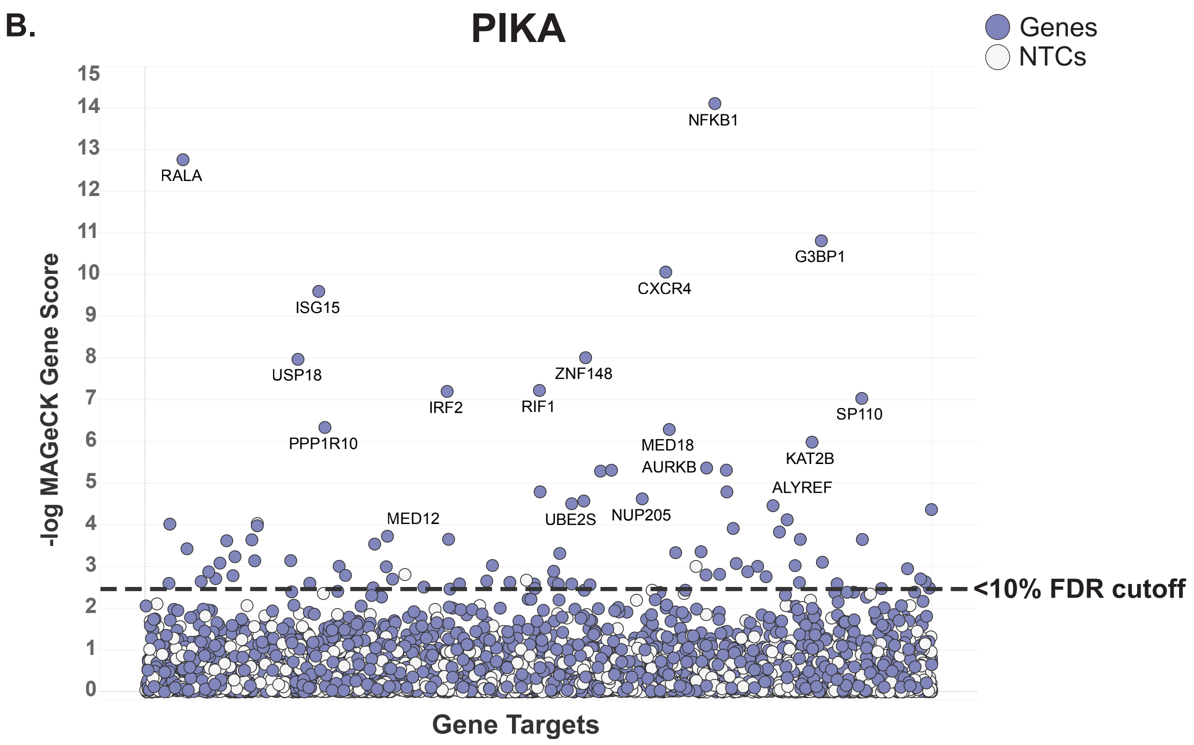

#### Supplemental Figure 2

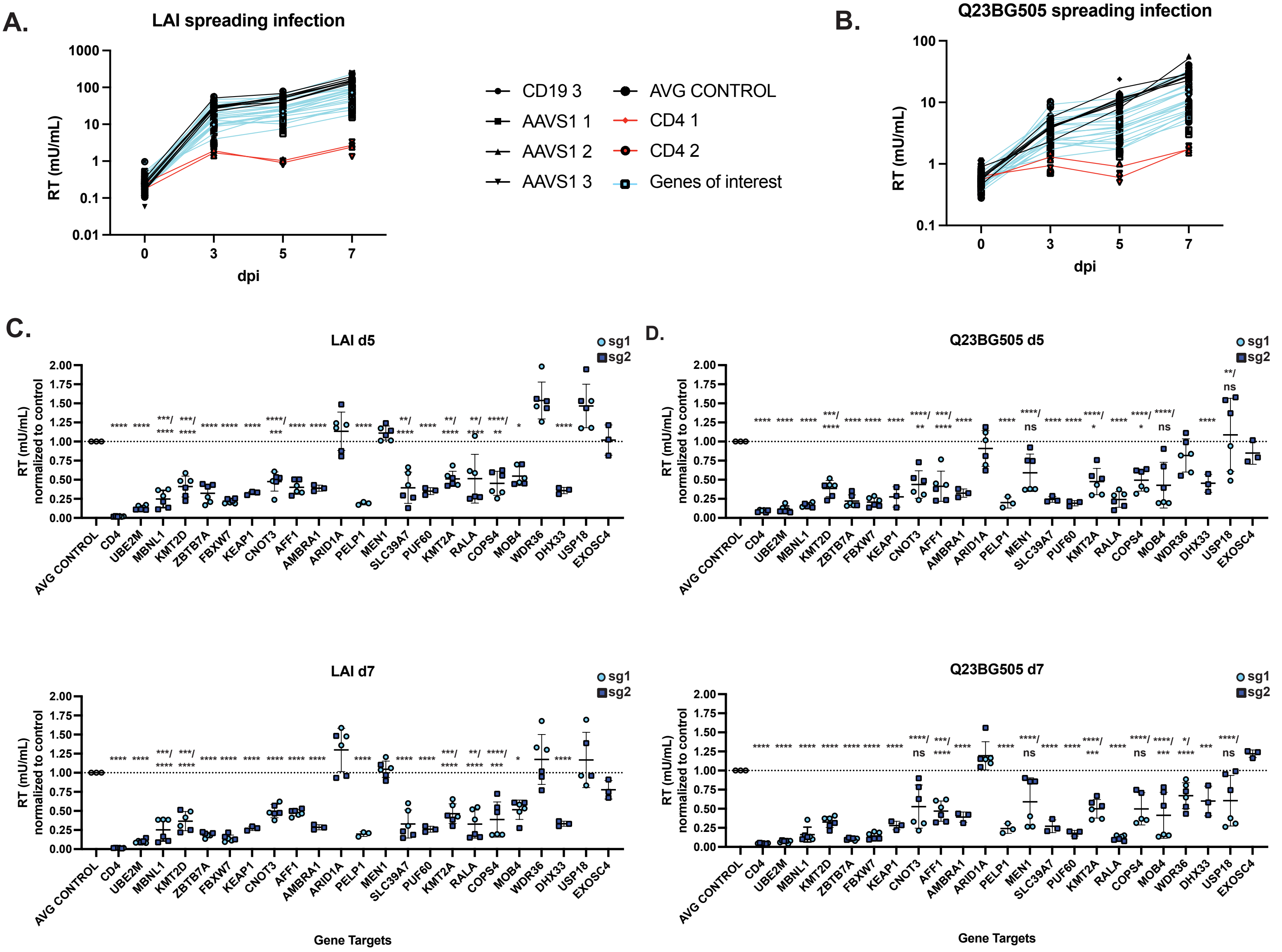

### Supplemental Figure 3

A.

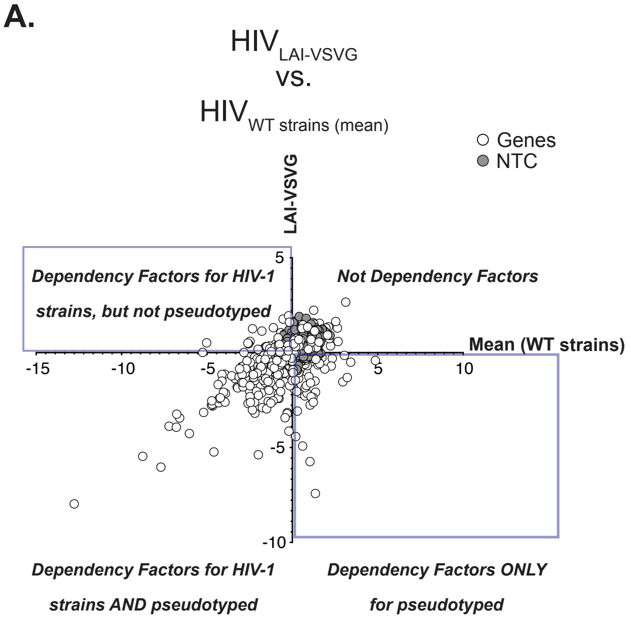


B.

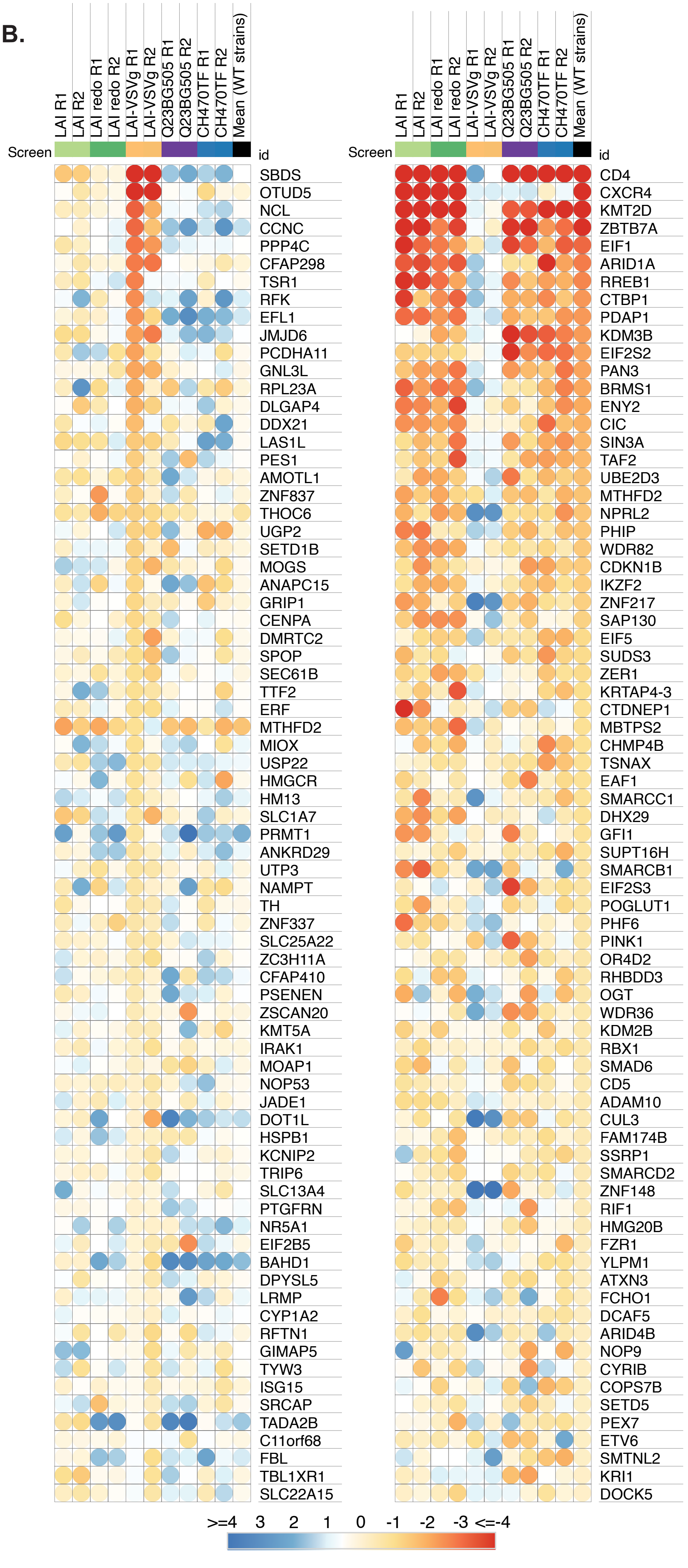

**Supplemental Figure 1.**

MAGeCK Gene analysis of the **(A)** Human Epigenome/Epigenetic library (HuEpi) or **(B)** Packageable ISG Knockout Assembly library (PIKA) screen showing the most depleted (dependency) genes in Jurkat T cells. X-Axis: randomly arrayed target genes. Y-Axis: -Log10 MAGeCK score for each host factors previously reported as dependency factors are shown in Pink. Purple represents the candidate genes. Genes scoring >10% FDR cutoff are those above the dashed line were included in the creation of the sublibrary HIVDEP. White = Non-targeting control (NTC) gene set generated in silico by randomly binning the 252 NTC sgRNA sequences into genes (six NTC guides per 841 genes) for HuEpi and 200 NTC sgRNA sequences into genes (eight NTC guides per 1,905 genes) for PIKA.

**Supplemental Figure 2.**

**(A-B)** Pooled ZapKO-Jurkat-CCR5 cells were generated by transducing with sgRNA/lentiCRISPRv2 lentiviruses targeting either positive control gene CD4, negative controls CD19 or AAVS1, or candidate dependency factor genes. One or two knockout lines per gene were generated using the highest scoring sgRNAs across each HIVDEP screen, selected in puromycin for >10 days to allow for gene knockout, followed by knockout efficiencies determination by ICE analysis (I) or infection with either LAI or Q23BG505 (MOI = 0.15). Viral supernatants were collected at days 0, 3, 5, and 7 to assess overall effect on replication kinetics via reverse transcriptase activity at each timepoint. Representative graphs were included in the main text. A and B represent the genes in the other batch. Y axis = Reverse Transcriptase milliUnits/ mL. **(C, D)** Reverse Transcriptase activity at the d5 and d7 timepoints were normalized to the mean of the control cell lines (CD19 and AAVS1 knockout lines) to combine batch 1 and 2 for each virus. Both knockout lines per gene are displayed: sgRNA 1 is shown as cyan circles and sgRNA 2 is shown as dark blue squares. For statistical analysis, all conditions are compared to the mean of the control cell lines (displayed as AVG CONTROLS). One way Anova; Tukey's multiple corrections test, p-value = ns > 0.05, <0.05 = *, =<0.05 = **, =<0.001 = ***, =<0.0001 = ****.

**Supplemental Figure 3.**

Correlation matrix and extended heatmaps of entry specific host factors.

**(A)** Correlation matrix of LAI-VSV-G screen z scores vs. mean of the wild-type strains, LAI, LAI redo, Q23BG505, and CH470TF screen z scores per each gene. **(B)** Entire heatmap of z scores arranged by the mean of the wild-type strains, from most to least (top 75).

**(C)** Entire heatmap of z scores arranged by the mean of the VSV-G-pseudotyped LAI, from most to least (top 75).

**Supplemental Table 1. MAGeCK gene scores for each screen**

**Supplemental Table 2. Complete list of the HIVDEP genes and sgRNA sequences**

**Supplemental Table 3. Z scores for each HIVDEP screen and Literature Review for top genes** (sorted by the mean of all stains excluding VSV-G pseudotyped LAI).

**Supplemental Table 4. Oligos/ primers used for CRISPR KO and ICE analysis**
